# Ongoing transposition in cell culture reveals the phylogeny of diverse *Drosophila* S2 sub-lines

**DOI:** 10.1101/2021.12.08.471819

**Authors:** Shunhua Han, Guilherme Dias, Preston J. Basting, Michael G. Nelson, Sanjai Patel, Mar Marzo, Casey M. Bergman

**Affiliations:** Institute of Bioinformatics, University of Georgia, Athens, GA, USA, 30602; Department of Genetics, University of Georgia, Athens, GA, USA, 30602; Faculty of Life Sciences, University of Manchester, Manchester, United Kingdom, M13 9PT

**Keywords:** *Drosophila*, transposable element, copy number variation, genome evolution, cell culture

## Abstract

Cultured cells are widely used in molecular biology despite poor understanding of how cell line genomes change *in vitro* over time. Previous work has shown that *Drosophila* cultured cells have a higher transposable element (TE) content than whole flies, but whether this increase in TE content resulted from an initial burst of transposition during cell line establishment or ongoing transposition in cell culture remains unclear. Here we sequence the genomes of 25 sub-lines of *Drosophila* S2 cells and show that TE insertions provide abundant markers for the phylogenetic reconstruction of diverse sub-lines in a model animal cell culture system. Analysis of DNA copy number evolution across S2 sub-lines revealed dramatically different patterns of genome organization that support the overall evolutionary history reconstructed using TE insertions. Analysis of TE insertion site occupancy and ancestral states support a model of ongoing transposition dominated by episodic activity of a small number of retrotransposon families. Our work demonstrates that substantial genome evolution occurs during long-term *Drosophila* cell culture, which may impact the reproducibility of experiments that do not control for sub-line identity.

## Introduction

Animal cell lines play vital roles in biology by providing an abundant source of material to study molecular processes and as cellular factories to express important biomolecules. Like all living systems, animal cell lines undergo genomic changes during routine propagation *in vitro* (Ruddle *et al*. 1958), leading to genetic diversity across time and laboratories that can lead to irreproducible research outcomes (Hughes *et al*. 2007). Despite the current emphasis on reducing sources of irreproducibility in biological research, relatively little attention has been paid to understand the pattern and process of *in vitro* evolution that leads to genomic diversity among sub-lines of long-term metazoan cell cultures (Junakovic *et al*. 1988; Di Franco *et al*. 1992; Ben-David *et al*. 2018; Liu *et al*. 2019), or how to identify and minimize the impact of such diversity (Hughes *et al*. 2007; Ben-David *et al*. 2018). Establishing general rules for cell culture genome evolution and mitigating its influence will likely require analysis of multiple cell lines from many different species since the pattern and process of genome evolution *in vivo* is known to vary across taxa (Lynch 2007).

Early studies in the model insect *Drosophila melanogaster* showed a high abundance of multiple transposable element (TE) families in cell lines relative to the genomes of whole flies (Potter *et al*. 1979; Ilyin *et al*. 1980). More recently, analysis of whole genome sequence (WGS) data revealed between ∼ 800 to ∼ 3000 non-reference TE insertions in different *Drosophila* cell lines, with LTR retrotransposons making up the bulk of these new insertions (Rahman *et al*. 2015). Proliferation of TEs in *Drosophila* cultured cell genomes could be explained by a burst of transposition during initial establishment of cell lines, by ongoing TE insertion during routine cell culture, or a combination of both processes (Echalier 1997). Di Franco *et al*. (1992) contrasted the stability of TE profiles among sub-lines of one of the oldest *Drosophila* cell lines (Kc) (Junakovic *et al*. 1988) with elevated TE abundance in a newly-established cell line (inb-c) and concluded that the increased TE abundance in *Drosophila* cell lines resulted from an initial burst of transposition during the establishment of a new cell line, with relative stasis thereafter. However, comparison of old and new cultures from different cell lines is not a definitive test of whether ongoing TE proliferation occurs during routine culture because of differences in the founder genotypes and cell type of independently established cell lines. More recently, Sytnikova *et al*. (2014) provided evidence for transposition after initial cell line establishment in *Drosophila* by showing an increase in abundance of the *ZAM* element in a continuously cultured sub-line of the OSS cell line (OSS_C) relative to a putative frozen progenitor sub-line (OSS_E). More recent work by Han *et al*. (2021) revealed that the early version of the OSS reported in Sytnikova *et al*. (2014) (OSS_E) is actually a misidentified version of a related cell line (OSC) and thus it is unclear if the *ZAM* activation in OSS occurred during or after the establishment of the OSS lineage. Documenting whether ongoing transposition in cell culture occurs is important since this process can lead to genomic variation among sub-lines that could impact functional studies and, more practically, provide useful markers for cell line identification and reconstruction of cell line evolutionary history (Han *et al*. 2021; Mariyappa *et al*. 2021).

Here we contribute to the understanding of genome evolution during long-term animal cell culture using a large sample of sub-lines of *Drosophila* Schneider Line 2 (S2) cells, one of the most widely-used non-mammalian cell culture systems. S2 cells were established from embryonic tissue of an unmarked stock of Oregon-R flies in December 1969 (Schneider 1972) and are likely to be derived from macrophage-like hemocytes (Schneider 1972; Echalier 1997). Two other cell lines, S1 (August 1969) and S3 (February 1970), were derived from the same ancestral fly stock (Schneider 1972) and can serve as outgroups to analyze evolution in the S2 lineage. Since their establishment, S2 cells have been distributed widely and grown more extensively than S1 or S3 cells (Lee *et al*. 2014). Many different sub-lines of S2 cells have been established by labs in the *Drosophila* community, some of which have been donated back to the *Drosophila* Genomics Resource Center (DGRC) for maintenance and distribution. In general, the provenance and relationships among sub-lines of S2 cells are unknown, as is the extent of their genomic or phenotypic diversity. At least one sub-type of S2 cells, called S2R+ (for S2 receptor plus), is known to have distinct phenotypes from other S2 cell lines such as expressing the Dfrizzled-1 and Dfrizzled-2 membrane proteins and having the desirable property of being more adherent to surfaces in tissue culture (Yanagawa *et al*. 1998). In addition to their ubiquity and diversity, S2 cells are a good model to study genome evolution in animal cell culture because of their relatively small genome size, which permits cost-effective whole-genome sequencing, and the wealth of prior biological knowledge in *D. melanogaster*.

In this study, we report new WGS data for 25 sub-lines of S2 cells as well as the outgroup S1 and S3 cell lines. We analyze these data together with public WGS samples for S2R+ and mbn2 (recently shown by Han *et al*. (2021) to be a misidentified lineage of S2) and demonstrate that TE insertions provide abundant markers to reconstruct the evolutionary history of S2 sub-lines. These data reveal that publicly available S2 sublines form a monophyletic group defined by two major clades (A and B), and suggest that misidentification of available S2 cultures by other *Drosophila* cell lines is limited. We also show that genome-wide copy number profiles support the major phylogenetic relationships among S2 sub-lines inferred using TE profiles. Using TE site occupancy and ancestral states, we infer that TE insertion has occurred on all internal branches of the S2 phylogeny, but that only a small subset of *D. melanogaster* TE families have proliferated during S2 evolution, most of which are retrotransposons that do not encode a retroviral envelope (*env*) gene. Together, these results support the conclusions that TE insertions provide useful markers of S2 sub-line identity and genome organization and that TE proliferation in *Drosophila* somatic cell culture is primarily driven by an ongoing, episodic, cell-autonomous process that does not involve deregulation of global transpositional control mechanisms.

## Materials and Methods

### Genome sequencing

We sequenced the genomes of 29 samples of S1, S2, or S3 cells to understand the genomic diversity and evolutionary relationships of publicly available sub-lines of S2 cells. Frozen stocks for each of these 29 samples were ordered from the *Drosophila* Genomics Resource Center (DGRC), American Type Culture Collection (ATCC), Deutsche Sammlung von Mikroorganismen und Zellkulturen (DSMZ), and Thermo Fisher. DNA was prepared directly from thawed samples without further culturing. Stock or catalogue numbers for these publicly available cell lines can be found in Table S1. Cells were defrosted and 250μl of the cell suspension was aliquoted and spun down for 5 min at 300g. The supernatant was discarded and the DNA from the cell pellet was extracted using the Qiagen DNeasy Blood & Tissue Kit (Cat. No. 69504). DNA preps were done in three batches, each of which contained an independent sample of S2-DRSC (DGRC-181) to identify any potential sample swaps and to assess the reproducibility of phylogenetic clustering based on TE profiles. The triplicate samples of S2-DRSC were from the same freeze of this cell sub-line performed by DGRC (Daniel Mariyappa, personal communication). Illumina sequencing libraries were generated using the Nextera DNA sample preparation kit (Cat. No. FC-121-1030), AMPure XP beads were then used to purify and remove fragments *<*100bp, and libraries were normalized and pooled prior to being sequenced on an Illumina HiSeq 2500 flow cell using a 101bp paired-end layout.

In addition, we analyzed public WGS data for a sample of S2R+ (unpublished results; G. Dias, S. Han, P. Basting, R. Viswanatha, N. Perrimon, and C.M. Bergman) and three samples of mbn2, a cell line which was recently shown to be a misidentified lineage of S2 cells (Han *et al*. 2021). A summary of the sequence data analyzed for each of the 33 samples in this study can be found in Table S1.

### Prediction of non-reference TE insertions

Non-reference TE insertions were detected in each sample using trimmed paired fastq sequences as input for the TEMP (Zhuang *et al*. 2014) module in McClintock (v2.0) (Nelson *et al*. 2017). We used TEMP to predict non-reference TEs based on previous results showing TEMP predictions are the least dependent on coverage and read length relative to other component methods in McClintock (Han *et al*. 2021). By default, McClintock filters predictions made by TEMP by requiring at least one read support on both sides of insertion and at least 10% TE allele frequency. The major sequences (chr2L, chr2R, chr3L, chr3R, chr4, chrM, chrY, and chrX) from the *D. melanogaster* dm6 assembly were used as a reference genome (Hoskins *et al*. 2015). The TE library used for McClintock analysis was a slightly modified version of the Berkeley *Drosophila* Genome Project canonical TE dataset described in Sackton *et al*. (2009) (https://github.com/bergmanlab/transposons/blob/master/releases/D_mel_transposon_sequence_set_v10.2.fa).

Genome-wide non-reference TE predictions generated by McClintock were filtered to exclude TEs in low recombination regions using boundaries defined by Cridland *et al*. (2013) lifted over to dm6 coordinates, as in Han *et al*. (2021). Our analysis was restricted to normal recombination regions since low recombination regions have high reference TE content which reduces the ability to predict non-reference TE insertions (Bergman *et al*. 2006; Manee *et al*. 2018). Low recombination regions included in our analyses were defined as chrX:405967–20928973, chr2L:200000–20100000, chr2R:6412495–25112477, chr3L:100000– 21906900, chr3R:4774278–31974278. We also excluded *INE-1* family from the subsequent analysis since this family has been reported to be inactive in *Drosophila* for millions of years (Singh and Petrov 2004; Wang *et al*. 2007). Filtered non-reference TE predictions were then clustered across genomic coordinates and samples. TEs predicted in different samples in the same cluster were required to directly overlap and be on the same strand. Clustered non-reference TE predictions were then filtered to exclude low-quality predictions using the same criteria as in Han *et al*. (2021). Briefly, non-reference TE loci with a single TE family per locus and one prediction per sample were retained.

### Phylogenetic analysis of cell sub-line samples using TE insertion profiles

Genome-wide non-reference TE predictions were then converted to a binary presence/absence matrix as input for phylogenetic analysis. Phylogenetic trees of cell sub-lines were built using Dollo parsimony in PAUP (v4.0a168) (Swofford 2003). Phylogenetic analysis was performed using heuristic searches with 50 replicates. A hypothetical ancestor carrying the assumed ancestral state (absence) for each locus was included as root in the analysis (Batzer and Deininger 2002; Han *et al*. 2021). “DescribeTrees chgList=yes” option was used to assign character state changes to all branches in the tree. Finally, node bootstrap support for the most parsimonious tree was computed by integrating 100 replicates generated by PAUP using SumTrees (v4.5.1) (Sukumaran and Holder 2010).

### Copy number analysis of cell sub-line samples

BAM files generated by McClintock were used to generate copy number profiles for non-overlapping windows of the dm6 genome using Control-FREEC (v11.6) (Boeva *et al*. 2012). 10 kb windows were used for Control-FREEC analyses unless specified otherwise. Windows with less than 85% mappability were excluded from the analysis based on mappability tracks generated by GEM (v1.315 beta) (Derrien *et al*. 2012). Baseline ploidy was set to diploid for S1 and tetraploid for all other samples, according to ploidy levels for S1, S2, S2R+, S3, and mbn2 cells estimated by Lee *et al*. (2014). The minimum and maximum expected value of the GC content was set to be 0.3 and 0.45, respectively.

## Results

### Genome-wide TE profiles reveal the evolutionary relationships among Schneider cell sub-lines

Previously, we showed that genome-wide TE profiles can be used to uniquely identify *Drosophila* cell lines and provide insight into the evolutionary history of clonally-evolving sub-lines derived from the same cell line (Han *et al*. 2021). Here, we propose that TE profiles can also be used to infer the currently unknown evolutionary relationships for a large panel of diverse sub-lines originating from a widely-used animal cell line, *Drosophila* S2 cells. We generated paired-end Illumina WGS data for a panel of 25 *Drosophila* S2 sub-lines from multiple lab origins (Table S1), including triplicate samples of one sub-line (S2-DRSC) to act as an internal control, and for the S1 and S3 cell lines that were derived from the same ancestral fly stock (Oregon-R) as the S2 lineage (Schneider 1972). In our analysis, we also included a S2R+ sub-line from the *Drosophila* RNAi Screening Center (DRSC) with publicly available WGS data from a forthcoming study (unpublished results; G. Dias, S. Han, P. Basting, R. Viswanatha, N. Perrimon, and C.M. Bergman) and three mbn2 cell sub-line samples from Han *et al*. (2021) (Table S1). mbn2 cells were originally reported to have a distinct origin (Gateff *et al*. 1980), but recent genomic analysis has shown that currently-circulating mbn2 cells are a mis-indentified lineage of S2 cells (Han *et al*. 2021), although it remains unknown to which lineage mbn2 cells are most closely related. Using TEMP (Zhuang *et al*. 2014), we predicted between 655 and 2924 non-reference TE insertions in the euchromatic regions of these Schneider cell line samples (Table S2). Each sample had a unique profile of non-reference TE insertions (File S1).

We performed phylogenetic analysis using genome-wide TE profiles of all Schneider cell line samples using the Dollo parsimony approach (Han *et al*. 2021). This approach fits the assumptions of the homoplasy-free nature of TE insertions (Shedlock and Okada 2000; Salem *et al*. 2003; Xing *et al*. 2005; Platt *et al*. 2015; Lammers *et al*. 2017, 2019) while also accommodating the false negative TE predictions inherent to short-read-based TE detection methods (Nelson *et al*. 2017; Rishishwar *et al*. 2017; Vendrell-Mir *et al*. 2019). The most parsimonious tree revealed several expected patterns that suggest using TE profiles to infer the evolutionary relationship among Schneider cell lines is reliable (Figure 1A; File S2). First, most internal nodes have high bootstrap support. All weakly-supported nodes are close to the terminal taxa, which presumably is due to the lack of phylogenetically-informative TE insertions that differentiate very closely related sub-lines or sample replicates. Second, using a hypothetical ancestor representing the state without any non-reference insertions to root the tree, S1 and S3 cell lines were independently reconstructed as outgroups for the S2 sub-lines in the phylogeny, as expected based on their independent origin from the same ancestral fly stock (Schneider 1972). Third, replicate samples of S2-DRSC cluster as nearest taxa and form a monophyletic clade with 100% bootstrap support. Fourth, all samples from S2R+, which are sub-lines of S2 with unique phenotypic characteristics (Yanagawa *et al*. 1998), form a monophyletic clade with 100% bootstrap support. Finally, all mbn2 sub-lines form a monophyletic clade with 100% bootstrap support embedded within a monophyletic clade of S2 sub-lines that itself has 100% bootstrap support. These results suggest that TE profiles can be used to reliably infer the evolutionary relationship among diverse sub-lines of a widely-used animal cell line, and that there is no evidence for any S2 sub-lines in our dataset being a misidentified non-S2 *Drosophila* cell lines.

**Figure 1.**
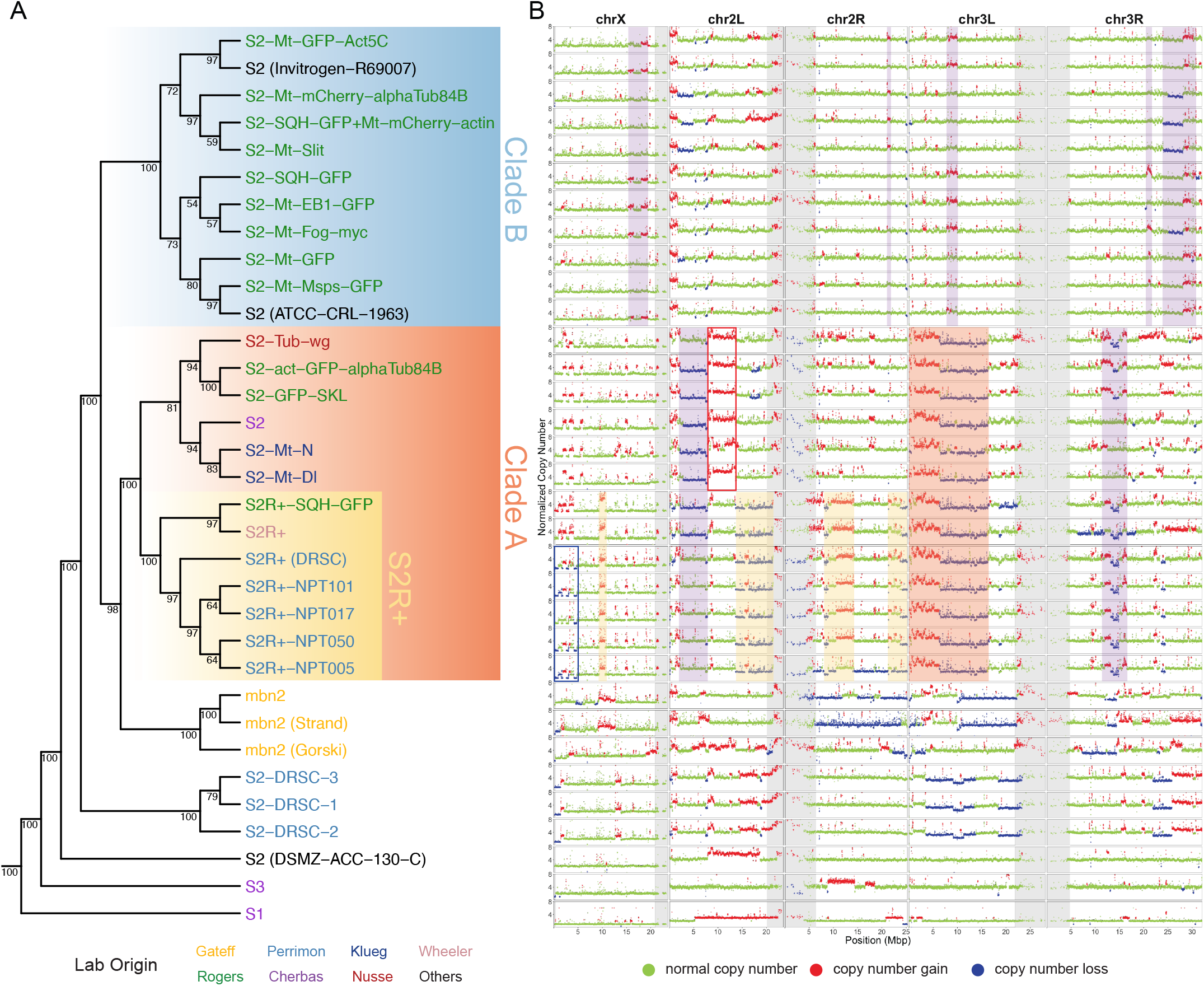
TE and CNV profiles reveal the evolutionary relationship among S2 sub-lines. (A) Dollo parsimony tree including a panel of 26 S2 sub-lines with diverse lab origins, two S1 and S3 sub-lines to serve as outgroups in the phylogeny, and three mbn2 sub-lines that were inferred to be misidentified S2 lines by Han *et al*. (2021). Replicate samples for S2-DRSC were also included. The phylogeny was constructed using genome-wide non-reference TE insertions predicted by TEMP (Zhuang *et al*. 2014). Percentage bootstrap support was annotated below each node. *Drosophila* Genomics Resource Center (DGRC) cell line names are used as taxa labels. Samples obtained from other sources are labeled in the format of “cell line name (source name)”. Taxa labels were colorized based on original labs in which cell sub-lines were developed. (B) Copy number profiles for samples included in panel A separated by chromosome arms. Each data point represents normalized copy number (ratio*ploidy) for a given 10kb window estimated by Control-FREEC (Boeva *et al*. 2012). Data points for each window are colorized by CNV status (red: CNV gain; green: no CNV; blue: CNV loss), which are based on the comparison between normalized copy number computed by Control-FREEC and baseline ploidy estimated by Lee *et al*. (2014). Red shading indicates CNVs that are exclusively shared by all S2 sub-lines in Clade A. Yellow shading indicates CNVs that are exclusively shared by S2R+ sub-lines. The red box in chromosome X represents CNVs that are exclusively shared by all S2 sub-lines in Clade A that are not S2R+. The blue box in chromosome arm 2L represents CNVs that are exclusively shared by S2R+ sub-lines from the Perrimon lab. Purple shading indicates CNVs that are exclusively and shared by a subset of S2 sub-lines within Clade A or Clade B. Low recombination regions are shaded in grey.

The phylogeny of Schneider cell lines built using TE profiles revealed a major split in the history of S2 cell line evolution, resulting in two sister lineages which we labelled as “Clade A” and “Clade B” (Figure 1). Clade A is comprised of one subclade containing all seven S2R+ sub-lines and another sub-clade containing six S2 sub-lines, one of which is the canonical S2 subline distributed by DGRC (DGRC-6). Clade B is comprised of 11 S2 sub-lines including sub-lines from Invitrogen and ATCC. The presence of S2 sub-lines in both clade A and clade B, but the presence of S2R+ sub-lines in clade B, implies that the S2 cell line designation is paraphyletic (i.e., some S2 sub-lines are more closely related to S2R+ than to other S2 sub-lines). In some cases, Schneider cell lines from the same lab cluster together (e.g. S2R+ sub-lines from the Perrimon lab and S2 sub-lines from the Klueg lab, respectively). However, S2 sub-lines from the Rogers lab were placed in different major clades of the S2 phylogeny (three S2-sub-lines in Clade A, nine S2-sub-lines in Clade B, Figure 1), demonstrating that the same lab can use divergent sub-lines of S2 from different major clades that have potentially different genome organization (see below).

The majority of S2 sub-lines we surveyed in this study were placed within Clade A and Clade B based on their TE profiles. However, two S2 sub-lines, S2-DRSC and S2 (DSMZ-ACC-130-C), were independently placed as outgroups for the two major clades of S2, suggesting that they are highly divergent S2 lineages. S2-DRSC is routinely used for RNAi screens at the *Drosophila* RNAi Screening Center (DRSC) and was recently donated to DGRC. Its relationship to the canonical S2 sub-line from DGRC (i.e., DGRC-6) was previously not known. Our results suggest that S2-DRSC and S2 (DGRC-6) are not closely related sub-lines, which could explain the phenotypic and functional differences between these two sub-lines reported in previous studies (Cherbas *et al*. 2011; Wen *et al*. 2014; Lee *et al*. 2014; Lee and Oliver 2015).

mbn2 sub-lines cluster in a monophyletic clade that is sister to Clade A (98% bootstrap support) but is clearly contained within a monophyletic lineage containing all S2 samples. This observation is consistent with previous results reported by Han *et al*. (2021) proposing that mbn2 is a misidentified S2 lineage. Han *et al*. (2021) showed that mbn2 clusters with S2-DRSC before clustering with S2R+. However, our results showed that the mbn2 clade clusters Clade A (containing S2R+ sub-lines) before clustering with S2-DRSC. We interpret this discrepancy as being caused by the sparse sampling and use of low coverage sequencing data for S2 and S2R+ from the modENCODE project in the previous study (Han *et al*. 2021), which led to insufficient signal to infer the evolutionary relationship of the mbn2 clade within S2 sub-line diversity.

### Genome-wide copy number profiles correlate with history of S2 sub-lines

To further investigate potential genomic heterogeneity among Schneider cell lines and cross-validate our phylogenetic reconstruction based on TE profiles, we generated copy number profiles for all samples in our dataset (Figure 1B) using Control-FREEC (Boeva *et al*. 2012). Two patterns in the copy number profiles generated suggested that our approach to characterize segmental variation in our cell sub-lines was robust. First, we observed a high concordance in copy number profiles for replicate samples of S2-DRSC (Figure 1B). Second, copy number profiles we generated using our new data for S1, S2R+, S2-DRSC, and S3 are broadly consistent with profiles for these cell lines using data generated by the modENCODE project reported previously in Lee *et al*. (2014) (Figure S2).

Copy number profiles for S2 sub-lines revealed a substantial amount of segmental copy number variants (CNVs) among different clades in the S2 phylogeny (Figure 1B). The major Clades A and B have distinct patterns of CNV variation, with S2 sublines in Clade A having many CNVs, while sub-lines in Clade B have very few CNVs throughout their genomes (Figure 1B). CNVs that are exclusively shared by sub-lines in Clade A but not present in clade B are readily apparent, such as the ∼ 15Mbp copy number gains and losses on chromosome arm 3L (Figure 1B, red shading). The two main sub-clades within Clade A are also distinguished by sub-clade-specific CNVs: several copy number gains and losses on chromosome X, arm 2L, and arm 2R are exclusively shared by all S2R+ sub-lines (Figure 1B, yellow shading), while a ∼5Mbp copy number gain on chromosome arm 2L is exclusively shared by non-S2R+ sub-lines (Figure 1B, red box). Within the S2R+ clade, there are also copy number losses in the distal regions of chromosome X that are exclusively shared by S2R+ sub-lines from the Perrimon lab (Figure 1B, blue box). Furthermore, S2-DRSC and S2 (DSMZ-ACC-130-C) have distinct copy number profiles that differ from other S2 sub-lines in Clade A and Clade B (Figure 1B), supporting the inference based on TE profiles that these are divergent S2 lineages. Finally, CNV profiles for mbn2 samples have distinct copy number profiles that differ from all other S2 sub-lines, consistent with the interpretation that mbn2 cells are a divergent lineage of S2. In addition, we note that the abundance and diversity of CNVs in mbn2 sub-lines resembles the CNV diversity observed for S2 sub-lines in Clade A (Figure 1B), the major S2 clade which the mbn2 is inferred to be most closely related to based on TE profiles.

We also observed some examples where reversals of CNVs may have arisen by somatic recombination or aneuploidy. For example, S2R+, S2R+-SQH-GFP, and most S2 sub-lines in Clade A (except S2-Tub-wg) share a ∼ 5Mbp copy number loss in chromosome arm 2L (Figure 1B). This pattern could be explained by a segmental deletion event occurring in the common ancestor of sub-lines in Clade A, followed by reversals of the deletion in S2-Tub-wg and in the common ancestor of S2R+ sub-lines from Perrimon lab through somatic recombination (Figure 1B). In addition, a copy number loss on the entire chromosome arm 2R can be observed for S2R+-NPT005 but not for other S2R+ sub-lines, which can be explained by a whole-arm aneuploidy event. Overall, these results suggest that copy number changes contribute to substantial diversity in genome organization among S2 sub-lines and that shared patterns of CNVs are broadly consistent with the evolutionary relationships among S2 sub-lines inferred from TE profiles (Figure 1A).

### Evidence for ongoing transposition during long-term S2 cell culture

In the absence of secondary events such as segmental deletion, ancestral non-reference TE insertions from the original fly strain or that arose during cell line establishment will be clonally-inherited by all descendant sub-lines. Ancestral insertions in regions without copy number loss should not provide any phylogenetic signal, and thus a simple model of TE proliferation during cell line establishment with no subsequent genome evolution cannot jointly explain (i) the overall increase in TE abundance and (ii) phylogenetically-informative nature of TE insertions in S2 cells. Two other contrasting models can however account for both features of the TE landscape in S2 genomes. Under the “Early transposition and subsequent deletion” model (Figure 2A), the increase in TE abundance is caused by a massive proliferation of TEs during cell line establishment, with subsequent copy number loss events shared by descendent cell lines indirectly explaining the phylogenetic signal of genome-wide TE profiles. Under the “Ongoing transposition in cell culture” model (Figure 2A), it is not necessary to invoke any TE proliferation during cell line establishment, and both the overall increase in TE abundance and phylogenetic signal of TE profiles result from the ongoing accumulation of TE insertions during routine cell culture that are inherited by descendent cell lines.

**Figure 2.**
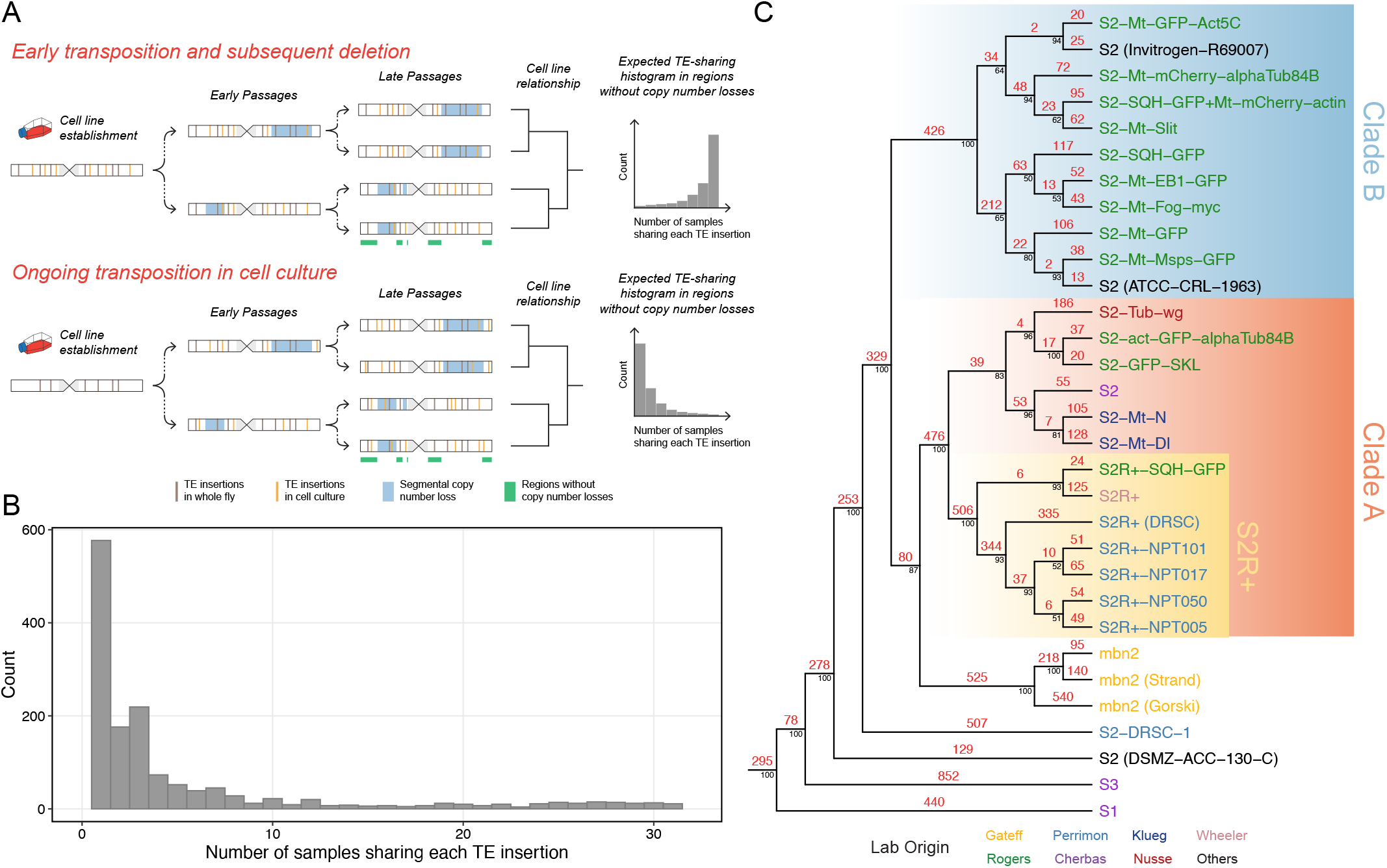
TE profiles suggest ongoing transposition in S2 cell culture. (A) Two hypotheses that could explain the mode of TE amplification in *Drosophila* S2 cell culture and how the resulting TE profiles could help infer the relationship among different cell sublines. Note that the schematic models represent genome-wide TE distributions combining all haplotypes. Therefore, given that S2 cells are tetraploid (Lee *et al*. 2014), a copy number loss event that occurred in one haplotype should only wipe out some TEs that are heterozygous in the affected region. (B) Histogram shows the distribution of the number of *Drosophila* S2 sub-line samples that share each TE insertion in regions of chromosome X without major shared copy number losses (chrX:500000-20928973). (C) Dollo parsimony tree including 26 *Drosophila* S2 sub-lines constructed using non-reference TE predictions made by TEMP (Zhuang *et al*. 2014). Samples from S1, S3, and mbn2 cell lines were also included. The number of TE insertions estimated using ancestral state reconstruction were annotated in red above each branch. Percentage bootstrap support was annotated in black below each node. *Drosophila* Genomics Resource Center (DGRC) cell line names are used as taxa labels. Samples obtained from other sources are labeled in the format of “cell line name (source name)”. Taxa labels were colorized based on original labs in which cell sub-lines were developed.

These alternative models can be distinguished by analyzing TE profiles in regions of the genome without copy number loss events. In regions without shared copy-number-loss events, the “Early transposition and subsequent deletion” model predicts that TE insertions will be shared by the majority of sub-lines and that TE profiles will not have strong phylogenetic signal to infer the evolutionary history of S2 sub-lines. In contrast, the “Ongoing transposition in cell culture” model predicts that few TEs will be shared by all sub-lines in regions without shared copy-number-loss events, and that TE profiles in these regions will be able to reconstruct evolutionary history of S2 sub-lines in a similar manner as genome-wide TE profiles. To test these alternative models, we analyzed TE profiles in a ∼ 15Mbp region in chromosome X that does not include significant copy number loss across all S2 sub-lines we surveyed (Figure S1B, purple shading). Our analysis revealed that the majority of TE insertions in regions of the X chromosome without shared copy number loss events are exclusive to one or a subset of S2 sub-line samples (Figure 2B). Phylogenetic analysis of non-reference TE insertions in the same region of chromosome X generated a most parsimonious tree that has the same major topological features as the one built from genome-wide TE profiles (Figure S1A). Together, these results provide evidence against the “Early transposition and subsequent deletion” model and suggest that the genomewide TE profiles used to infer evolutionary relationship of S2 sub-lines are contributed mainly by ongoing lineage-specific transposition during cell culture.

### A subset of LTR retrotransposon families have episodically inserted during S2 cell line history

To gain additional insights into the dynamics of TE activity during the history of S2 cell line evolution, we mapped TE insertions on the phylogeny of *Drosophila* S2 sub-lines using ancestral state reconstruction based on the most parsimonious scenario of TE gain and loss under the Dollo model (Batzer and Deininger 2002; Ray *et al*. 2006; Han *et al*. 2021) (Figure 2C). The Dollo model favors TE insertions to be gained once early in the phylogeny over parallel gains of TEs in different sub-lineages (Farris 1977) and is thus conservative with respect to the number of inferred transposition events on more terminal branches of the tree. The most parsimonious reconstruction of TE insertions mapped on the Schneider cell line phylogeny reveals a substantial number of TE insertions on branches at all depths in the phylogeny (Figure 2C). For example, we observe over 250 TE insertions on each ancestral branch that split the divergent S2 lineages S2-DRSC and S2 (DSMZ-ACC-130-C) from the major S2 clades, and more than 400 TE insertions on the ancestral branches leading to both major Clades A and B. Likewise, more than 500 TE insertions are mapped on the ancestral branch leading to the S2R+ clade. This pattern of abundant insertion on most major internal branches of the phylogeny provides further support to the “Ongoing transposition in cell culture” model.

We then aggregated inferred TE insertions on each branch by TE family to visualize branch- and family-specific TE insertion profiles. This analysis revealed that only a subset of 125 recognized TE families in *D. melanogaster* contribute to the high transpositional activity in S2 cell culture (Figure 3B; File S3). The top ten TE families with highest overall activities are all retrotransposons, including eight LTR retrotransposons (*blood, copia, 297, 3S18, 1731, diver, mdg1* and *17*.*6*) and two non-LTR retrotransposons (*jockey* and *Juan*). The majority of the most active TE families in S2 cells do not encode a retroviral *env* gene (8/10; 80%), with only the *297* and *17*.*6* Ty3/gypsy families having the potential to form infectious virus-like particles (Lerat and Capy 1999; Malik *et al*. 2000; Stefanov *et al*. 2012). This analysis also revealed that the pattern of TE family activity varies substantially on different branches of the S2 phylogeny (Figure 3). For example, families such as *17*.*6, 297*, and *1731* have relatively high activity in branches prior to the split of Clade A and B (branch 33-36; “early S2”) and in the early branches within Clade A and S2R+ (branch 48,49), but relatively low activity within Clade B. In contrast, families such as *jockey, blood*, and *3S18* have relatively low activity in “early S2” branches and relatively high activity across all branches within Clade A and B. We also observed TE family activity that is sub-line-specific, including the proliferation of *gtwin* that occurred only in S2-Mt-Dl (Figure 3), a sub-line of S2 that was transformed to express wild-type Delta from a Cu-inducible metallothionein promoter (FBtc0000152). Together, these results suggest that the increase in abundance of TEs during S2 cell culture is caused by a small subset of retrotransposon families, and that there have been episodic periods of family-specific transposition during the evolutionary history of S2 cells.

**Figure 3.**
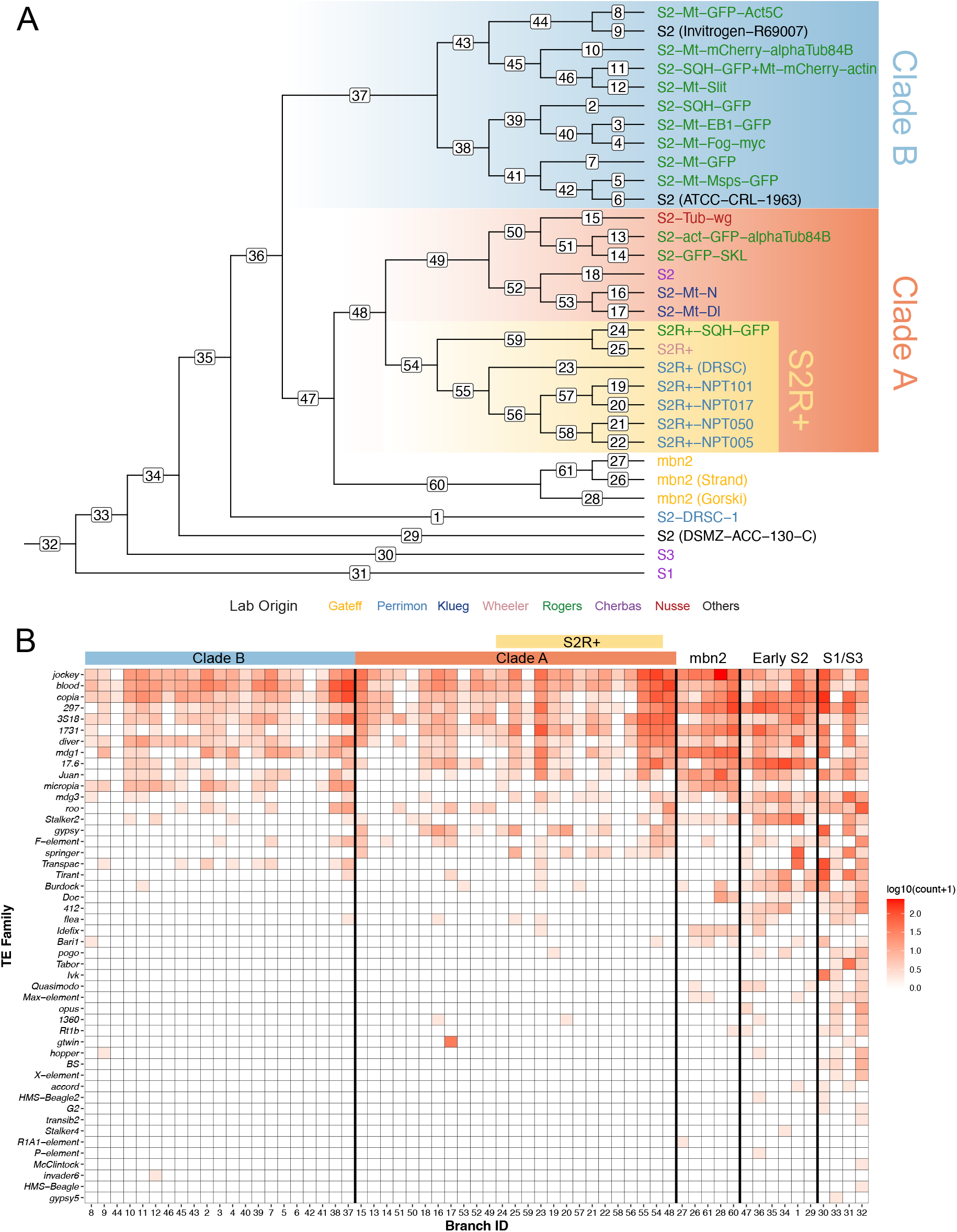
Ongoing transposition in *Drosophila* S2 culture is contributed by a small subset of LTR retrotransposon families. (A) Dollo parsimony tree including 26 *Drosophila* S2 sub-lines constructed using non-reference TE predictions made by TEMP (Zhuang *et al*. 2014). Samples from S1, S3, and mbn2 cell lines were also included. Taxa labels were colorized in the same way as Figure 1 and Figure 2C. Branch ID is annotated on each branch. (B) Heatmap showing the number of estimated family-specific TE insertions on each branch of the tree in panel A. The heatmap is colorized by log-transformed (log10(count+1)) number of gains per family per branch, sorted top to bottom by overall non-reference TE insertion gains per family across all branches, and sorted left to right into clades representing major clades of S2 phylogeny with major clade color codes indicated at the top of the heatmap.

## Discussion

Here we used genome-wide TE and copy number profiles to reveal the evolutionary relationships and genomic diversity among a large panel of diverse *Drosophila* S2 sub-lines. Our TE-based phylogenetic analysis showed that all S2 sub-lines sampled form a single monophyletic clade that is an ingroup to the expected outgroup S1 and S3 cell lines. This result suggests that no S2 sub-line in our dataset is a misidentified non-S2 *Drosophila* cell line, and implies relatively low rates of cross-contamination in the community between S2 cells and other *Drosophila* cell lines. Our results also revealed two major clades of S2 sub-lines that are supported by copy number profiles. One major clade that we labeled as “Clade A” includes all S2R+ sub-lines and several S2 sub-lines. This clade is characterized by substantial copy number changes across the autosomes. The other major clade we labeled as “Clade B” and includes only S2 sub-lines with mostly euploid genomes. These results imply that the “S2” subline designation is paraphyletic and that there can be substantial genomic heterogeneity among sub-lines labeled as S2. We also found that some S2 sub-lines originating from the same lab were reconstructed in different major clades of S2, providing evidence that heterogeneity in S2 genome content has the potential to influence experimental results within a single laboratory. We note that since we do not have information about the number of passages leading to each sample in our dataset, we cannot quantitatively relate how TE insertion or copy number changes occur as a function of evolutionary time. Thus, differences in genomic variability among Clades A and B may simply reflect the number of passages rather than intrinsic differences in genome stability. Future mutation accumulation experiments would be needed to estimate rates of transposition and copy number evolution in S2 cell culture and could help date the divergence time among major branches of the S2 tree.

Our phylogeny of S2 sub-lines also clarifies the origin of S2R+ cells, a lineage whose increased adherence to tissue culture surfaces has led to its use in nearly 600 primary publications (FBtc0000150). S2R+ cells were first reported by Yanagawa *et al*. (1998) who showed that S2R+ cells are responsive to Wingless signaling and expressed the Wingless receptors Dfrizzled-1 and Dfrizzled-2, in contrast to S2 cells from the Nusse lab (presumably represented by a Clade A sub-line like S2-Tub-wg). Yanagawa *et al*. (1998) report that the founding sub-line of the S2R+ lineage was obtained from Dr. Tadashi Miyake Lab, who stated that these cells were “were obtained directly from Dr. Schneider and stored frozen in his laboratory.” This reported history has led the DGRC to conclude that S2R+ cells are “more similar to the original line established in the Schneider laboratory than any of the other S2 isolates in our collection.” (https://dgrc.bio.indiana.edu/cells/S2Isolates). In contrast to this reported history, our results place the S2R+ lineage as a derived clade inside Clade A, rather than at the base of the S2 phylogeny as would be expected if S2R+ cells were a basal lineage that reflects the original state of all S2 sub-lines. Furthermore, our results indicate that the increased adherence and Wingless responsiveness of S2R+ cells are derived features, suggesting that they may have arisen as adaptations to propagation in cell culture. Further work will be necessary to understand the mechanisms that caused the *in vitro* evolution of these phenotypes, however preliminary analysis suggests that the gain of expression for Dfrizzled-1 and Dfrizzled-2 was not caused by increased copy number in the ancestor of S2R+ sub-lines, nor is the inferred lack of expression of these genes in other S2 isolates due to complete deletion of these loci (Figure S3).

Our phylogenetic hypothesis for the evolution of Schneider cell lines also allowed us to test competing models to explain the proliferation of TEs in *Drosophila* cell culture. Analysis of TE site occupancy in regions of the genome without shared copy number loss provided evidence against the “Early transposition and subsequent deletion” model while supporting the “Ongoing transposition in cell culture” model. Likewise, analysis of ancestral states provided additional evidence for the “Ongoing transposition in cell culture” model. One potential issue with our analysis of inferred TE ancestral states is the possibility of false-positive (FP) and false-negative (FN) non-reference TE predictions. In principle, a random FP prediction is unlikely to be shared by multiple cell samples and thus should only lead to a falsely reconstructed insertions on the terminal branches under the Dollo model. This suggests that the number of TE insertions reconstructed on the terminal branches of our trees may be overestimated. Conversely, a random FN would most likely lead to falsely reconstructed deletion on the terminal branch under the Dollo model. Thus, random FP and FN TE predictions should have a limited impact on our phylogenetic and ancestral state reconstruction analyses and thus not majorly affect the conclusion that there are substantial numbers of TE insertions on most internal branches of the tree, as expected under the “Ongoing transposition in cell culture” model.

Additionally, our ancestral state reconstruction analysis revealed that only a subset of TE families have high transpositional activity in S2 cell culture. Most active TE families in S2 cells are retrotransposons that do not encode a functional retroviral *env* gene and thus are not capable of infecting another cell, suggesting that TE proliferation in *Drosophila* cell culture is mainly a cell-autonomous process. Furthermore, the fact that we do not observe activation of all TE families suggests transposition in S2 is not due to global deregulation of all TEs but is caused by some form of family-specific regulation. Finally, our ancestral state reconstruction analysis revealed that transposition of active TE families in S2 culture is episodic. Some TE families such as *17*.*6, 297*, and *1731* have relatively higher activities in the early stage of S2 evolution, while other families such as *jockey, blood*, and *3S18* were more active within two major clades of S2. Arkhipova *et al*. (1995) provided two non-mutually exclusive hypotheses for proliferation of TEs in cell lines: 1) ongoing transposition is more easily tolerated in cultured cells and is no longer under strong negative selection as it is in the whole flies; or 2) there exist specific factors that control TE transposition, and their actions are altered significantly in cell culture. Our observation of family-specific, episodic TE activity during S2 cell line evolution favors changes in TE regulation over global relaxation of selection to explain TE proliferation in *Drosophila* cell culture. However, more work is needed to understand the mechanism by which TE copy number regulation is relaxed in a family-specific fashion in S2 cells and other *Drosophila* cell lines.

Overall, this study revealed ongoing somatic TE insertions and copy number changes as mechanisms for genome evolution in *Drosophila* S2 cell culture in the 50 years of its history since establishment (Schneider 1972). These results provide new insights into cell line genome evolution for a non-human metazoan species, and add to the genomic and phenotypic heterogeneities within cell culture that have been reported for the human HeLa cell line (Liu *et al*. 2019) and MCF-7 breast cancer cell lines (Ben-David *et al*. 2018). Together, these findings suggest that rapid genome evolution and sub-line heterogeneity are common features of animal cell lines evolving *in vitro*. Future work is needed to further characterize the rates and patterns of cell line genome evolution in a diversity of systems to better understand how *in vitro* genome evolution changes affect cell line phenotypes and functional outcomes.

## Data Availability

Raw sequencing data generated in our study is available in the SRA under BioProject PRJNA603568. Supplemental Material available at TBD. File S1 contains nonredundant BED files from McClintock runs using TEMP module on the dataset including 33 *Drosophila* cell line samples (reference TEs, *INE-1* insertions and TEs in low recombination regions excluded). File S2 contains clustered TE profiles in the format of binary presence/absence data matrix including 33 *Drosophila* cell line samples (reference TEs, *INE-1* insertions and TEs in low recombination regions excluded). File S3 includes data matrix of the number of non-reference TE insertion gain events per family on each branch of the most parsimonious tree used for the heatmap in Fig. 3B.

## Acknowledgements

We thank Stacey Holden and Andy Hayes (University of Manchester Genomic Technologies Core Facility) for assistance with Illumina library preparation and sequencing; Shan-Ho Tsai and Yecheng Huang (University of Georgia) for bioinformatics application support; and the Georgia Advanced Computing Resource Center (University of Georgia) for computing time. We thank members of the Bergman Lab (University of Manchester and University of Georgia), and the Dyer, Hall, Sweigart and White Labs (University of Georgia) for helpful suggestions throughout the project. This work was supported by Wellcome Trust Award 096602/B/11/Z (MGN), University of Georgia Research Education Award Traineeship (PJB), Human Frontier Science Program grant RGY0093/2012 (CMB), and the University of Georgia Research Foundation (CMB).

## Notes

### Competing Interest Statement

The authors have declared no competing interest.

