## Supplemental Figures and Tables for "Ongoing transposition in cell culture reveals the phylogeny of diverse *Drosophila* S2 sub-lines"

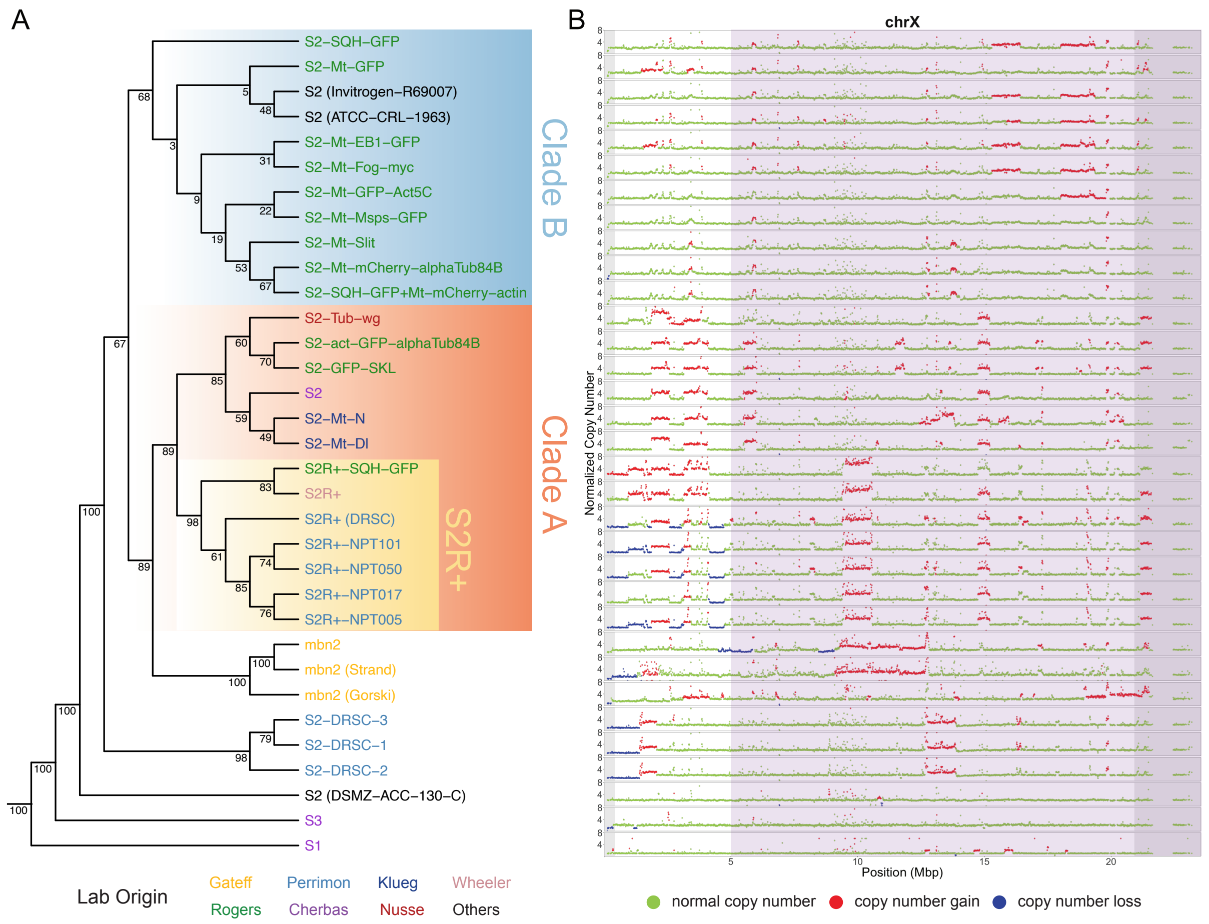


**Figure S1. Evolutionary relationship among S2 sub-lines inferred using TE profiles in regions without major shared copy number losses.** (A) Dollo parsimony tree including a panel of 26 *Drosophila* S2 sub-lines constructed using non-reference TE insertions predicted by TEMP (Zhuang *et al.*, 2014) in regions of chromosome X without major shared copy number losses (chrX:500000-20928973) indicated by purple shading. Samples from S1, S3, and mbn2 cell lines and replicate samples for S2-DRSC were also included. Percentage bootstrap support was annotated below each node. *Drosophila* Genomics Resource Center (DGRC) cell line names are used as taxa labels. Samples obtained from other sources are labeled in the format of “cell line name (source name)”. Taxa labels were colorized based on donor labs in which cell sub-lines were developed. (B) Copy number profiles of chromosome X for samples included in panel A. Each data point represents normalized copy number (ratio*ploidy) for a given 10 kb window estimated by Control-FREEC (Boeva *et al.*, 2012). Data points for each window are colorized by CNV status (red: CNV gain; green: no CNV; blue: CNV loss), which are based on the comparison between normalized copy number estimated by Control-FREEC and baseline ploidy estimated by Lee *et al.* (2014). Regions without major shared copy number losses among S2 sub-lines are shaded in purple. Low recombination regions are shaded in grey.

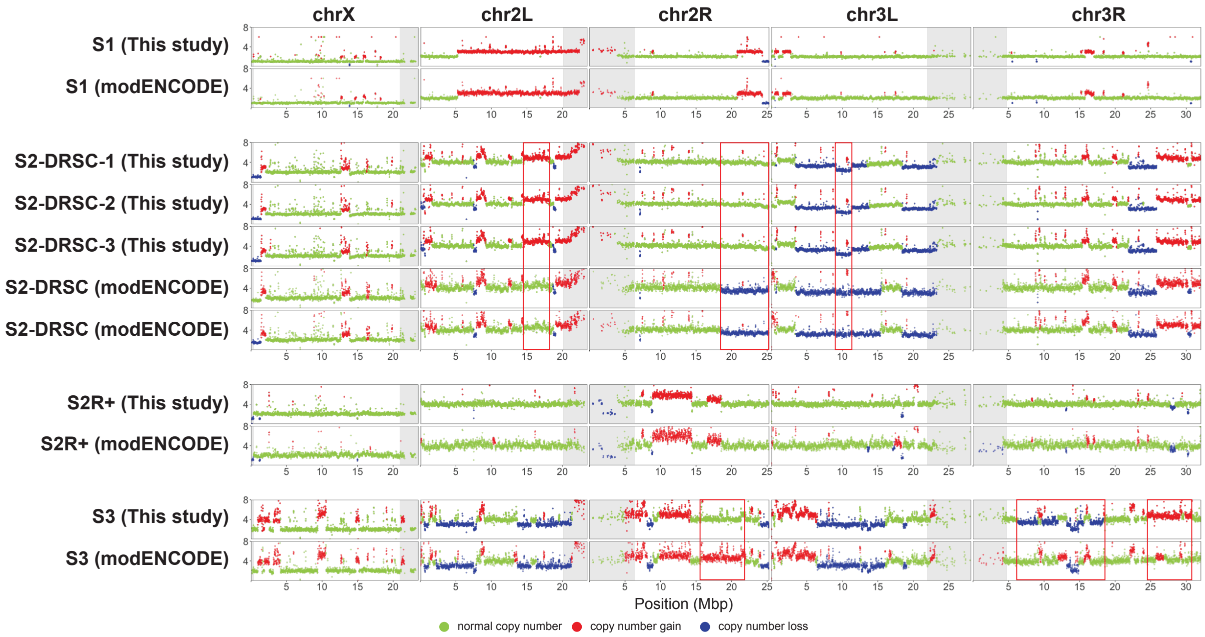


**Figure S2. Copy number profiles of four Schneider cell sub-line samples using data collected from this study and the modENCODE project.** Each data point represents normalized copy number (ratio*ploidy) for a given 10 kb window estimated by Control-FREEC (Boeva *et al.*, 2012). Data points for each window are colorized by CNV status (red: CNV gain; green: no CNV; blue: CNV loss), which are based on the comparison between normalized copy number estimated by Control-FREEC and baseline ploidy estimated by Lee *et al.* (2014). For each cell line sample, regions that exhibit differences in major CNV patterns between this study and the modENCODE project are marked in red boxes. Low recombination regions are shaded in grey.

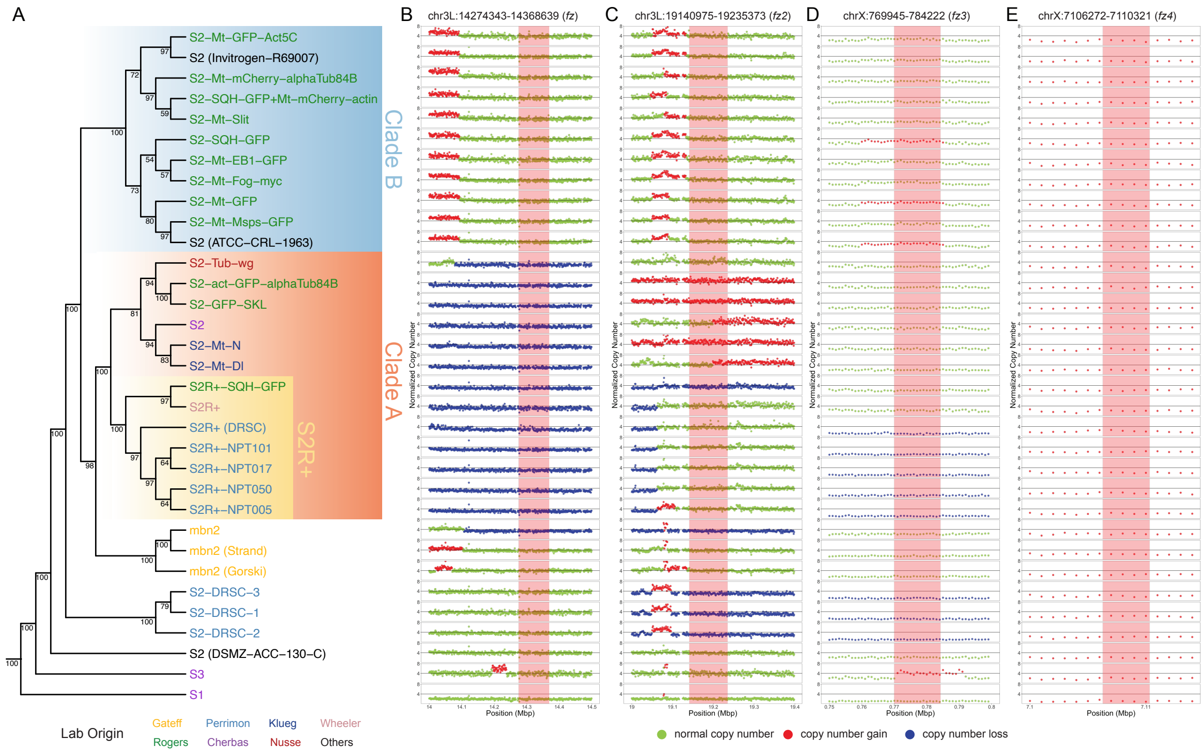


**Figure S3. Copy number profiles of four *frizzled* genes for Schneider cell sub-line samples.** (A) Dollo parsimony tree including a panel of 26 *Drosophila* S2 sub-lines constructed using non-reference TE predictions made by TEMP (Zhuang *et al.*, 2014). Samples from S1, S3, and mbn2 cell lines and replicate samples for S2-DRSC were also included. Percentage bootstrap support was annotated below each node. *Drosophila* Genomics Resource Center (DGRC) cell line names are used as taxa labels. Samples obtained from other sources are labeled in the format of "cell line name (source name)". Taxa labels were colorized based on donor labs in which cell sub-lines were developed. (B-E) Copy number profiles of *fz*, *fz2*, *fz3*, and *fz4* gene loci for samples included in panel A. Each data point represents normalized copy number (ratio*ploidy) for a given 1 kb window estimated by Control-FREEC (Boeva *et al.*, 2012). Data points for each window are colorized by CNV status (red: CNV gain; green: no CNV; blue: CNV loss), which are based on the comparison between normalized copy number estimated by Control-FREEC and baseline ploidy estimated by Lee *et al.* (2014). Red shading indicates *frizzled* gene regions.

**Table S1. Summary of 31 Schneider cell line samples analyzed in this study.** *Drosophila* Genomics Resource Center (DGRC) cell line names are given for all cell line samples except for samples obtained from other sources, which are labeled in the format of "cell line name (source name)". The "Lab Origin" represents the lab that originally created the sub-line. "Inferred ploidy" and "Inferred sex" represent the ploidy and sex of the cell line, respectively, estimated by Lee *et al.* (2014). "Read pairs" represents the number of paired-end reads for a given cell sub-line sample. "Coverage" represents the average mapped depth of coverage after quality and adaptor trimming. N.A. indicates that this information is not available.

| Cell line | DGRC ID | Lab origin | Inferred ploidy | Inferred sex | SRA | Read length | Read pairs | Coverage |
|---|---|---|---|---|---|---|---|---|
| mbn2 | DGRC-147 | Gateff | 4 | male | SRR13360020 | 151 | 55531647 | 109.61 |
| mbn2 (Gorski) | N.A. | Gateff | 4 | male | SRR13360019 | 151 | 63738692 | 126.11 |
| mbn2 (Strand) | N.A. | Gateff | 4 | male | SRR13360018 | 151 | 68440069 | 132.00 |
| S1 | DGRC-9 | Cherbas | 2 | male | SRR10981795 | 101 | 34904345 | 35.79 |
| S2 | DGRC-6 | Cherbas | 4 | male | SRR10981796 | 101 | 28189507 | 31.67 |
| S2 (ATCC-CRL-1963) | N.A. | Others | 4 | male | SRR10981814 | 101 | 50088154 | 47.26 |
| S2 (DSMZ-ACC-130-C) | N.A. | Others | 4 | male | SRR10981794 | 101 | 51683568 | 48.34 |
| S2 (Invitrogen-R69007) | N.A. | Others | 4 | male | SRR10981793 | 101 | 43038240 | 42.18 |
| S2-act-GFP-alphaTub84B | DGRC-170 | Rogers | 4 | male | SRR10981789 | 101 | 37705915 | 37.57 |
| S2-DRSC-1 | DGRC-181 | Perrimon | 4 | male | SRR10981786 | 101 | 31515040 | 34.76 |
| S2-DRSC-2 | DGRC-181 | Perrimon | 4 | male | SRR10981812 | 101 | 49916928 | 51.87 |
| S2-DRSC-3 | DGRC-181 | Perrimon | 4 | male | SRR10981811 | 101 | 50084326 | 49.18 |
| S2-GFP-SKL | DGRC-197 | Rogers | 4 | male | SRR10981805 | 101 | 47302631 | 48.96 |
| S2-Mt-Dl | DGRC-152 | Klueg | 4 | male | SRR10981802 | 101 | 37297961 | 38.88 |
| S2-Mt-EB1-GFP | DGRC-171 | Rogers | 4 | male | SRR10981788 | 101 | 34454428 | 39.07 |
| S2-Mt-Fog-myc | DGRC-218 | Rogers | 4 | male | SRR10981803 | 101 | 38085006 | 40.59 |
| S2-Mt-GFP | DGRC-194 | Rogers | 4 | male | SRR10981808 | 101 | 41882171 | 43.14 |
| S2-Mt-GFP-Act5C | DGRC-169 | Rogers | 4 | male | SRR10981790 | 101 | 44383445 | 46.85 |
| S2-Mt-mCherry-alphaTub84B | DGRC-195 | Rogers | 4 | male | SRR10981807 | 101 | 44960162 | 44.06 |
| S2-Mt-Msps-GFP | DGRC-206 | Rogers | 4 | male | SRR10981804 | 101 | 47561052 | 48.99 |
| S2-Mt-N | DGRC-154 | Klueg | 4 | male | SRR10981792 | 101 | 40733228 | 43.91 |
| S2-Mt-Slit | DGRC-192 | Rogers | 4 | male | SRR10981810 | 101 | 41033908 | 41.86 |
| S2-SQH-GFP | DGRC-172 | Rogers | 4 | male | SRR10981787 | 101 | 26690953 | 29.73 |
| S2-SQH-GFP+Mt-mCherry-actin | DGRC-193 | Rogers | 4 | male | SRR10981809 | 101 | 42525580 | 41.74 |
| S2-Tub-wg | DGRC-165 | Nusse | 4 | male | SRR10981791 | 101 | 41826501 | 43.40 |
| S2R+ | DGRC-150 | Wheeler | 4 | male | SRR10981813 | 101 | 20056094 | 23.23 |
| S2R+ (DRSC) | N.A. | Perrimon | 4 | male | SRR11000336 | 151 | 47640215 | 82.65 |
| S2R+-NPT005 | DGRC-229 | Perrimon | 4 | male | SRR10981801 | 101 | 45413880 | 45.17 |
| S2R+-NPT017 | DGRC-230 | Perrimon | 4 | male | SRR10981800 | 101 | 34459982 | 35.27 |
| S2R+-NPT050 | DGRC-231 | Perrimon | 4 | male | SRR10981799 | 101 | 29162882 | 28.27 |
| S2R+-NPT101 | DGRC-232 | Perrimon | 4 | male | SRR10981798 | 101 | 43114190 | 43.26 |
| S2R+-SQH-GFP | DGRC-196 | Rogers | 4 | male | SRR10981806 | 101 | 49206117 | 47.08 |
| S3 | DGRC-5 | Cherbas | 4 | male | SRR10981797 | 101 | 23764412 | 27.52 |

**Table S2. Number of non-reference TE predictions made by TEMP for 31 Schneider sub-lineage samples.** Numbers of non-reference TE insertion predictions made by TEMP are based on default McClintock (Nelson *et al.*, 2017) settings. *INE-1* and non-reference TE insertion predictions in low recombination regions were excluded. N.A. indicates that this information is not available.

| Cell line | DGRC ID | SRA | # of TEs |
|---|---|---|---:|
| mbn2 | DGRC-147 | SRR13360020 | 1934 |
| mbn2 (Gorski) | N.A. | SRR13360019 | 2195 |
| mbn2 (Strand) | N.A. | SRR13360018 | 1980 |
| S1 | DGRC-9 | SRR10981795 | 743 |
| S2 | DGRC-6 | SRR10981796 | 1268 |
| S2 (ATCC-CRL-1963) | N.A. | SRR10981814 | 847 |
| S2 (DSMZ-ACC-130-C) | N.A. | SRR10981794 | 655 |
| S2 (Invitrogen-R69007) | N.A. | SRR10981793 | 845 |
| S2-act-GFP-alphaTub84B | DGRC-170 | SRR10981789 | 975 |
| S2-DRSC-1 | DGRC-181 | SRR10981786 | 1281 |
| S2-DRSC-2 | DGRC-181 | SRR10981812 | 1083 |
| S2-DRSC-3 | DGRC-181 | SRR10981811 | 1057 |
| S2-GFP-SKL | DGRC-197 | SRR10981805 | 866 |
| S2-Mt-Dl | DGRC-152 | SRR10981802 | 1263 |
| S2-Mt-EB1-GFP | DGRC-171 | SRR10981788 | 1230 |
| S2-Mt-Fog-myc | DGRC-218 | SRR10981803 | 1293 |
| S2-Mt-GFP | DGRC-194 | SRR10981808 | 1323 |
| S2-Mt-GFP-Act5C | DGRC-169 | SRR10981790 | 804 |
| S2-Mt-mCherry-alphaTub84B | DGRC-195 | SRR10981807 | 1032 |
| S2-Mt-Msps-GFP | DGRC-206 | SRR10981804 | 1053 |
| S2-Mt-N | DGRC-154 | SRR10981792 | 1134 |
| S2-Mt-Slit | DGRC-192 | SRR10981810 | 1038 |
| S2-SQH-GFP | DGRC-172 | SRR10981787 | 1533 |
| S2-SQH-GFP+Mt-mCherry-actin | DGRC-193 | SRR10981809 | 994 |
| S2-Tub-wg | DGRC-165 | SRR10981791 | 1210 |
| S2R+ | DGRC-150 | SRR10981813 | 1820 |
| S2R+ (DRSC) | N.A. | SRR11000336 | 2924 |
| S2R+-NPT005 | DGRC-229 | SRR10981801 | 1435 |
| S2R+-NPT017 | DGRC-230 | SRR10981800 | 1607 |
| S2R+-NPT050 | DGRC-231 | SRR10981799 | 1281 |
| S2R+-NPT101 | DGRC-232 | SRR10981798 | 1494 |
| S2R+-SQH-GFP | DGRC-196 | SRR10981806 | 1300 |
| S3 | DGRC-5 | SRR10981797 | 1204 |
